# Mechanisms of selection for the control of action in *Drosophila melanogaster*

**DOI:** 10.1101/296962

**Authors:** Giovanni Frighetto, Mauro A. Zordan, Umberto Castiello, Aram Megighian

**Affiliations:** Department of General Psychology, University of Padova, 35131, via Venezia 8, Padova, Italy.; Department of Biology, University of Padova, 35121, via G. Colombo 3, Padova, Italy.; Department of Biomedical Sciences, University of Padova, 35131, via U. Bassi 58/B, Padova, Italy.; Padova Neuroscience Center, University of Padova.

**Author notes:** All Authors partecipated equally to writing the manuscript. **Summary statement:** In this study we investigated adult fly locomotor behaviour in response to distracting stimuli during free walking. Kinematic data reveal an interesting phenomenon of motor re-orientation.

**Keywords:** Action selection, selective attention, motor re-orientation, model organism, fixation, visuo-motor integration, *Drosophila melanogaster*

## Abstract

In the last few years several studies have investigated the neural mechanisms underlying spatial orientation in *Drosophila melanogaster*. Convergent results suggest that this mechanism is associated with specific neural circuits located within the Central Complex (CC). Furthermore such circuits appear to be associated with visual attention, specifically with selective attention processes implicated in the control of action. Our aim was to understand how wild-type flies react to the abrupt appearance of a visual distractor during an ongoing locomotor action. Thus, we adapted the well-known ‘Buridan paradigm’, used to study walking behaviour in flies, so we could specifically address the mechanisms involved in action selection. We found that flies tended to react in one of two ways when confronted with a visual distractor during ongoing locomotion. Flies either: (i) committed to a new path situated midway between the original target and the distractor, consistent with a novelty effect; or (ii) remained on the original trajectory with a slight deviation in direction of the distractor. We believe that these results provide the first indication of how flies react, from the motor point of view, in a bi-stable context requiring the presence of selection-for-action mechanisms. Some considerations on the neural circuits underlying such behavioural responses are advanced.

## INTRODUCTION

Living organisms have evolved neural information processing systems to allow interaction with the environment so as to maximize the probability of survival and reproduction. To reach this goal, appropriate information about the environment has to be extracted by perceptual systems in a form that can be used to guide actions (Tipper et al., 1998; Castiello, 1999). Visual attention systems appear to operate by mapping out relevant perceptual aspects of the environment and translating them into appropriate action control parameters. *Drosophila melanogaster* also seems to employ such mechanisms, for instance, in order to avoid predator attacks, to prevent collisions with obstacles or to head efficiently towards salient visual stimuli (Card and Dickinson, 2008; van Breugel and Dickinson, 2012; Maimon et al., 2008). Therefore, it is reasonable to assume that the presence of efficient action selection mechanisms constitute an evolutionarily conserved characteristic (Strausfeld and Hirth, 2013; Grillner and Robertson, 2016). The putative neural substrate of an action selection system in flies is thought to be contained within a doughnut-shaped structure called the ellipsoid body (EB) (Fiore et al., 2015), which is part of a wider ensemble of modular neuropils involved in locomotor behaviour termed the central complex (CC), (Strauss and Heisenberg, 1993; Martin et al., 1999; Pfeiffer and Homberg, 2014). Recently, using a two-photon calcium imaging technique, it has been shown that a class of CC neurons termed E-PG neurons – having their dendritic tree in the EB and their axonal branches in the Protocerebral Bridge (PB) and Gall brain regions – are involved in tracking the orientation of a visual landmark and, to a lesser extent also the direction of body motion (Seelig and Jayaraman, 2015). The neurons of this circuit are arranged in a toroidal pattern, functionally subdividable into wedges. Each wedge responds to a particular direction of navigation through a mechanism involving a ring attractor dynamic model which explains how information concerning visual landmarks is integrated with self-motion in order to allow navigation (Turner-Evans et al., 2017; Heinze, 2017). Furthermore, this circuit is thought to be the neural centre for visual attention since it is characterized by a discrete single ‘bump’ of activity following the presentation of multiple visual stimuli (de Bivort and van Swinderen, 2016). This is reminiscent of a sort of attentional focus (Castiello and Umiltà, 1990; Castiello and Umiltà, 1992) and suggests a unified neurophysiological phenomenon which could form the basis of selection for the programming of locomotion direction.

Despite the above mentioned neurophysiological findings underlying landmark selection, little is known regarding the heading control in free moving adult flies. Horn and Wehner (1975)showed that walking flies faced with two stripes presented concomitantly and separated by an angular distance of less than 60 deg, preferred to move along the direction determined by the bisector of the angle between the two objects (Horn and Wehner, 1975). Conversely, when angles greater than 75 deg were considered, the flies showed a distribution of orientations with two maxima directed toward either of the two stripes. This behaviour has been described in terms of a superposition of two turning-tendency functions, which are phase shifted according to the angle subtended by the landmarks (Poggio and Reichardt, 1973; Horn and Wehner, 1975). In the light of recent findings, suggesting that the E-PG neurons operate according to ring attractor dynamics (Kim et al., 2017), it might be speculated that, in the case of Horn and Wehner’s experiment (1975), the ‘compass needle’ of the ring attractor points in a direction which is midway between the two landmarks. According to this idea, it has been observed that on some occasions E-PG activity transitioned from one offset to another relative to the two landmarks, indicating that this ambiguity may lead the fly to adopt an intermediate orientation (Seelig and Jayaraman, 2015). Thus, the turning tendency underlying fixation behaviour and the ring attractor model of the CC could be two sides of the same coin (Bahl et al., 2013; Seelig and Jayaraman, 2013).

With this in mind, we tested how the abrupt presentation of a visual stimulus to flies which are already engaged in locomotion (walking) toward a pre-existing visual target, would determine the activation of selection-for-action mechanisms which are then deployed in movement kinematics. To this end, we capitalized on an innate fly behaviour (i.e. unconditioned) in which flies continue to freely run back and forth between two opposing stripes inside a circular open arena (i.e. ‘Buridan’s paradigm’; Götz, 1980; Bülthoff et al., 1982; Strauss and Heisenberg, 1993; Strauss and Pichler, 1998). In our modified ‘Buridan paradigm’ a second stripe (with respect to the fly’s visual field) was presented while the fly was already moving towards the pre-existing stimulus. We hypothesize that the appearance of the extra stripe might determine three possible scenarios: i) if the presence of the second stripe does not alter the originally programmed direction of locomotion, then the fly’s movement should proceed in the direction of the first stripe, with no apparent changes along the path of the locomotion trajectory; ii) if the presence of the second stripe has a distracting effect, and therefore needs to be inhibited in order for the fly to proceed in the originally planned direction, then some evidence of this inhibitory process might be detectable in the form of slight perturbations in the locomotion trajectory; iii) if the presence of the second stripe determines the insurgence of an alternative motor program, which has the power to override the original one, then a dramatic change in direction toward the novel stripe should be evident. Surprisingly, the appearance of the novel target seemed to produce a tendency in the flies to turn towards a point midway between the two targets, as already shown by Horn and Wehner (1975). However, a more in depth analysis of the trajectories led us, in fact, to the identification of two alternative specific locomotor behaviours, namely that flies either: (i) committed to a new path situated midway between the original target and the distractor, consistent with a novelty effect; or (ii) presented a slight deviation of the original trajectory in the direction of the distractor. This in turn allows for interesting considerations regarding the nature of the selection-for-action mechanism in *Drosophila melanogaster*. In particular, the first type of response implies the abortion of the ‘old’ motor program in favour of a new one, while the second type of response suggests the deployment of an inhibitory mechanism operationalized in the form of slight trajectory changes.

## MATERIALS AND METHODS

### Animals

The experiments were performed on adult wild-type fruit flies *(Drosophila melanogaster*; Oregon-R strain). All flies were reared on standard cornmeal-sucrose-yeast medium at 22°C in a 12 h light/12 h dark cycle at 60% relative humidity. Fly crowding was also controlled (20-30 flies each vial) to avoid competition for food. Only individual 2-5 day-old male flies were used. Flies were kept in their food vials until the beginning of the experiment. Thus for the experiment flies were not starved nor were their wings clipped. All experiments were conducted between zeitgeber time 2 and 4 at room temperature 22-23°C.

### Experimental setup

To test how flies respond to the sudden appearance of new visual stimulus (distractor) during free walking toward a fixed visual stimulus (block) we employed a cylindrical led-emitting-diode (LED) modular display (Reiser and Dickinson, 2008) positioned around the fly (Fig. 1), and consisting in 48 (12 × 4) LED panels (each panel made by an 8 × 8 array of LEDs) (IO Rodeo Inc, Pasadena, CA, USA). A custom-designed transparent arena made of 3D-printed resin (iMaterialise HQ, Leuven, BE, EU) was placed within the cylindrical LED display. The cylindrical LED display and the transparent arena were in turn mounted on solid stainless steel brackets which were fixed to an aluminium breadboard (Thorlabs Inc, Newton, NJ, USA). The setup was thus positioned on an anti-vibration table, protected by a Faraday cage and covered with heavy black fabric. The arena (maximum height at the centre = 3.5 mm; diameter = 109 mm) was designed so as to i) confine flies in 2D space, ii) not allow the flies to reach the edge of the arena and iii) to impede flight by means of a glass ‘ceiling’ (Simon and Dickinson, 2010). The arena was backlit by an infrared (IR) LED array source (LIU850A, Thorlabs Inc, Newton, NJ, USA) and the IR light was diffused using paper diffuser films placed between the IR light source and the arena. A CCD camera (Chameleon 3, FLIR System Inc, Wilsonville, OR, USA) with 1288 × 964 pixel resolution, fitted with a 2.8-8 mm varifocal lens (Fujifilm, Tokyo, JP) and an 850 nm band pass filter (MidOpt Inc, Woodwork Lane Palatine, IL, USA) was mounted 36 cm above the arena in order to record fly activity. Videos of flies moving in the arena were recorded at 21 frames s^−1^, following selection of a 700 × 700 pixel region of interest which included the entire arena. In order to allow the experimenter to visually observe all events occurring within the arena (including whether visual patterns were being correctly displayed) an HD webcam (C310, Logitech, Lausanne, CH, EU) was also mounted alongside the infrared camera.

**Fig. 1.**
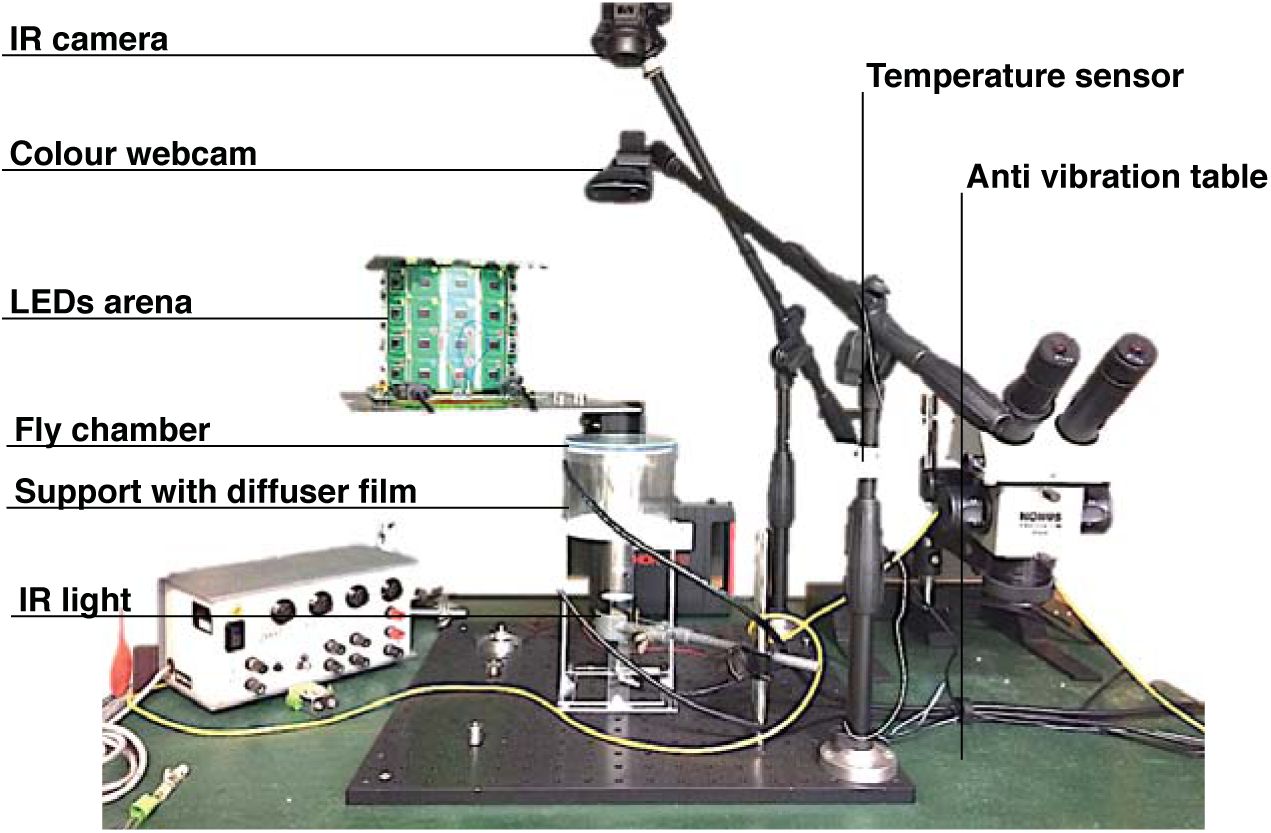
Experimental setup. Image showing the main components of the setup utilized in the experiment described in the paper.

### Procedure

Flies were individually loaded into the arena with a mouth aspirator and were left to adapt in complete darkness for at least 5 min. Individuals were then subjected to a ‘Buridan’s paradigm’, by illuminating two opposed bright stripes of 4 × 16 LEDs (width x height) each one covering 15 deg of the fly’s visual field when observed from the centre of the chamber. The classical interpretation of the phenomenon underlying this paradigm refers to the alternation between fixation and anti-fixation of attractive landmarks represented by black stripes on a bright background (Bülthoff et al., 1982). Apparently, bright stripes on a dark background show no difference in terms of fixation (Horn and Wehner, 1975; Seelig and Jayaraman, 2013). Preliminary experiments in our experimental setup showed a more robust response to the ‘Buridan’s paradigm’ in individuals tested with bright stripes on dark background, and, therefore, we decided to run our experiments with this setup. In our experiments, individual fly locomotion in ‘Buridan’s paradigm’, consisting in the fly continuously running to and fro between two opposing bright targets, was initially recorded for 200s (see Movie 1 in supplementary materials). Flies which did not exhibit this behaviour (i.e. remained still or roamed at random) were not further considered (Kain et al., 2012). This allowed to distinguish between flies that adopted a behaviour termed ‘quantum search action’ (i.e. a fixation and anti-fixation strategy) from those which did not manifest such behaviour. In other words, this procedure was aimed at selecting the ‘searcher’ phenotype considered for the following part of the experiment (Bülthoff et al., 1982). At the end of this selection phase, the behavioural task-proper was initiated. While the fly was still performing the ‘Buridan paradigm’, a second visual target (distractor) was presented the moment the fly crossed the virtual midline of the arena while moving between the two opposing bright stripes (a modified detour paradigm; Neuser et al., 2008). Therefore, our task consisted in a classical ‘Buridan paradigm’ performed under two alternative conditions. A distraction condition in which a single distracting-signal (chosen randomly among four alternative signals) was presented concomitantly with the ‘Buridan paradigm’ stimuli, whenever the individual crossed a virtual central window (27 mm width x 3.6 mm depth; see Fig. 2) along the chosen path. From this point on we shall refer to this condition as the ‘distractor’ condition. Distractors consisted in bright stripes of the same dimensions as the Buridan stripes (i.e. 15 deg of the fly’s visual field when viewed from the center of the arena). The distractors appeared randomly to the right or left of the fly at an angle of either 30 or 60 deg with respect to an ideal line connecting the opposing Buridan stripes. Each time a fly crossed the virtual central window, the distractor appeared for a 3 s period. During this period the two opposing Buridan stripes were always present. A ‘block’ condition, instead, consisted in the presentation of the Buridan stripes without any distractor. The experiment ended when the ‘block’ and the ‘distractor’ conditions had been presented seven times (with an average experiment lasting 30 min).

**Fig. 2.**
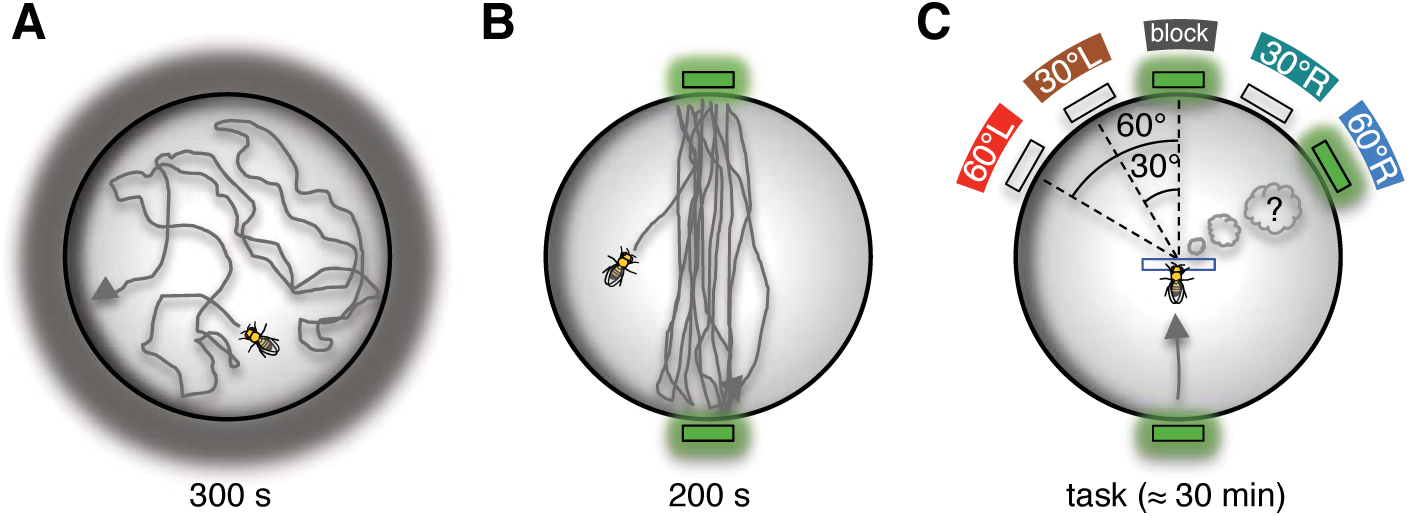
Experimental procedure. Cartoon showing the three phases involved in each experiment. (A) Acclimatization period in complete darkness for 300 s. (B) Two opposing bright green stripes were switched on and the behaviour was recorded for 200 s. (C) Behavioural task consisting in the random presentation of distracting visual stimuli (distractors) whenever the fly crossed a virtual central window (rectangle with blue borders). Behavioural task terminated when all five conditions were repeated seven times (about 30 min). Distractors are represented as: ‘block’ for no distraction, ‘30°R’ and ‘60°R’ for distraction at 30 or 60 deg on the right; ‘30°L’ and ‘60°L’ for distraction at 30 deg and 60 deg on the left.

### Software and management

The cylindrical LED display was controlled using available MATLAB (MathWork Inc, Natick, MA, USA) scripts (Reiser and Dickinson, 2008). The MATLAB Image Acquisition Toolbox was used to implement the system for video recording. Furthermore, in order to detect the position of the fly’s head in a specific spatial location (i.e. inside the virtual central window within the circular arena) and activate the necessary visual patterns on the LED panels accordingly, we implemented a system for real time tracking using the FAST (Features from Accelerated Segment Test) method (Rosten and Drummond, 2006) provided by the MATLAB Computer Vision System Toolbox. Online tracking analysis, video recording and control of the LED arena were integrated into a single custom GUI (Graphical Unit Interface), thus providing us with a unified software environment to manage all experimental variables. All the scheduled events involved in each experiment were automatically controlled by means of a custom script.

### Off-line tracking

To obtain a greater definition of the fly’s 2D position and body orientation we decided to track the fly’s trial recordings off-line using the CTRAX open source software (Branson et al., 2009). Errors occurring during tracking were fixed manually using appropriate available MATLAB scripts (CTRAX, FixErrors Toolbox) (Branson et al., 2009). Finally, other available MATLAB scripts (CTRAX, Behavioral Microarray Toolbox) were used to compute a suite of speed and acceleration properties (Branson et al., 2009).

### Data pre-processing

The files obtained following the off-line tracking analysis pipeline described above were transformed into .txt files, and imported into the R software (R Development Core Team, 2017) environment for data pre-processing and an initial exploratory analysis by means of custom scripts. For the trajectory analysis, only data from tracks in which single flies were directed towards the target were selected (i.e. all tracks in the opposite direction were removed). The minimum track length considered for analysis was 9 mm (i.e. 50 pixels; spatial resolution was 5.5 pixels per mm). Using this data frame (see Table 1 and Table 2) we performed track-centering. This operation proved necessary due to the fact that, in order to trigger the appearance of the distractor and to start the video recording, the flies had to cross a virtual central window within the circular arena. Given the dimensions of this virtual window, the tracks showed scattered starting-points along the x-axis (width of the window), depending upon the point at which the fly entered the virtual window. Therefore, since we were interested in evaluating the deviation of the fly locomotion paths caused by the different distractors and since the body orientations of the flies were uniform among conditions (Fig. 3A), we centered the starting point of all tracks at x = 0. Due to the limited depth of the triggering window the starting y values appeared to be more homogeneously distributed among the experimental conditions (Fig. 3B). Nonetheless, for uniformity, tracks were also centered at y = 0.

**Table 1.** Velocity and Distance with respect to the experimental condition

| Condition | No. trajectories | Velocity (mm s<sup>-1</sup>)<br>($\mu \pm \sigma$) | Distance (mm)<br>($\mu \pm \sigma$) |
| --- | --- | --- | --- |
| block | 123 | 18.83±13.62 | 46.64±13.32 |
| 30°R | 137 | 18.43±13.60 | 48.23±12.74 |
| 30°L | 135 | 18.50±13.87 | 48.80±13.09 |
| 60°R | 127 | 18.19±13.68 | 47.41±14.33 |
| 60°L | 131 | 18.44±13.54 | 49.88±14.39 |

**Table 2.** Number of trials by fly

| Fly ID | No.<br>trajectories | Fly ID | No.<br>trajectories | Fly ID | No.<br>trajectories |
| --- | --- | --- | --- | --- | --- |
| Fly_1 | 32 | Fly_8 | 30 | Fly_15 | 32 |
| Fly_2 | 34 | Fly_9 | 34 | Fly_16 | 29 |
| Fly_3 | 27 | Fly_10 | 32 | Fly_17 | 30 |
| Fly_4 | 30 | Fly_11 | 33 | Fly_18 | 27 |
| Fly_5 | 31 | Fly_12 | 34 | Fly_19 | 32 |
| Fly_6 | 33 | Fly_13 | 30 | Fly_20 | 31 |
| Fly_7 | 34 | Fly_14 | 31 | Fly_21 | 27 |

**Fig. 3.**
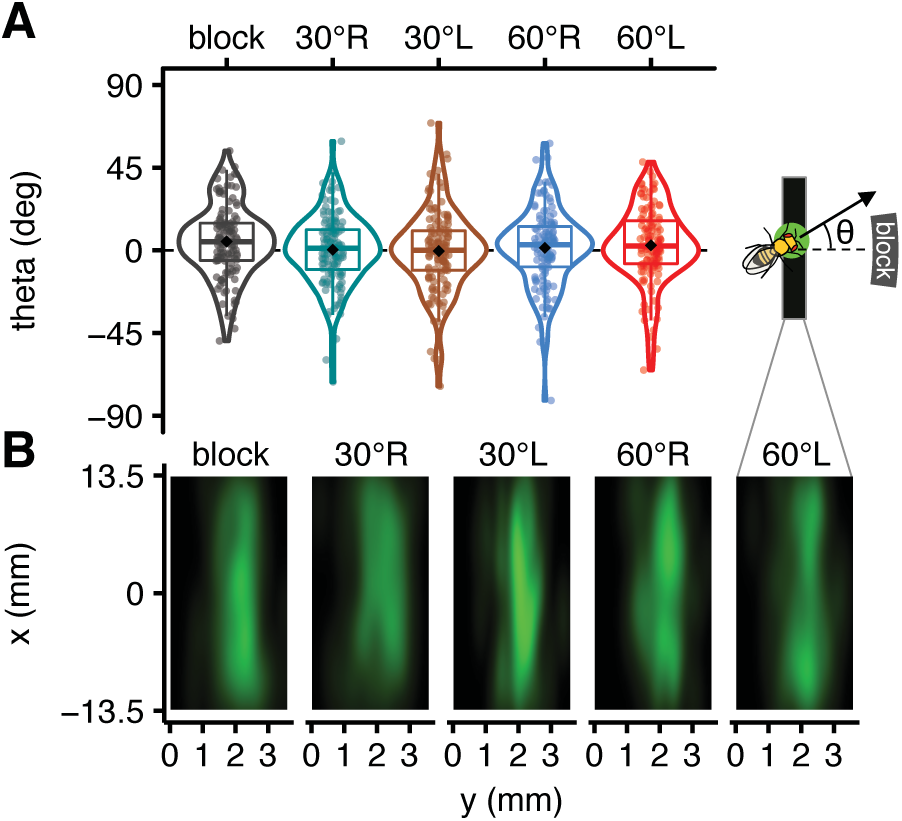
Data inspection and trajectories centering. (A) Box-violin plot (i.e. box plot plus data distribution) of flies orientations (theta; see inset) in degrees at the exact moment of distractor presentation. Plots show that flies orientations when faced with the trials do not differ consistently among different conditions and are approximately normally distributed. Colour coding: black correspond to the absence of distractors (block); green to distractor at 30 deg on the right side (30°R); brown to distractor at 30 deg on the left (30°L); blue to distractor at 60 deg on the right (60°R) and red to distractor at 60 deg on the left (60°L). Negative theta values refer to right-hand turns, while the positive ones to left-hand turns. The box-violin plot shows two measures of central tendency, the median in the box plot, and the mean of the data represented by the black square dot. (B) Heat map showing a density plot of all flies positions in the virtual rectangle when the distractor is presented, x and y-axis are in mm.

### Statistical approach

In order to understand how the presence of distractors explained the orientation and the trajectories taken by the flies we tested a series of Linear Mixed Effects (LME) models using the R package *lme4* (Bates et al., 2014). We used LME because such models allow to adjust estimates for repeated sampling (i.e. more than one observation arises from the same fly) and for imbalance in sampling (i.e. some flies are sampled more than others). LME also allow to take into account the experimental variation (i.e. variation among flies or among other groupings within the data) and to avoid the harmful effects of averaging, since this tends to remove variation (McElreath, 2016). Subsequently, the LMEs were compared in order to select the best model (i.e. the best fit to the data). For model selection we used the Bayesian Information Criterion (BIC) also known as the Schwarz information criterion or Schwarz’s BIC (Schwarz, 1978), an index that measures the efficiency of the model in terms of data forecasting. Since BIC tends to favour models with fewer parameters, we further conducted a Bayes Factor (BF) analysis with a method based on the multivariate generalizations of Cauchy priors (JZS method, see Liang et al., 2008) using the R package *BayesFactor* (Morey and Rouder, 2015). We used this parameterization because it allows BF to have excellent statistical properties independently of the phenomenon under study (a method also known as ‘objective Bayesian’, see Wagenmakers, 2007). The Bayes Factor expresses the ratio between the plausibility of observed data under M1 (our best model) and the plausibility of observed data under M0 (the null model). We compared different models, and the one with the highest Bayes Factor (greatest plausibility) was selected. With the *BayesFactor* package, which inherits the MCMC (Markov Chain Monte Carlo) sampling algorithm from the R package *coda* (Plummer et al., 2006), we were also able to compute the posterior distributions of parameters (with 10000 iterations). This approach to estimating parameters enabled us to take maximum advantage of LME modelling, which provided the direct probability of an effect (i.e. posterior probability) as well as the computation of the evidence for the results. Analysis of variance (ANOVA) and non-parametric Mann-Whitney-Wilcoxon tests were also used, under the null hypothesis that the sample distributions belonged to the same population.

## RESULTS

### Orientation effects

As a first step we investigated the body orientation adopted by the flies at the time the second visual stimulus (i.e. the distractor) was presented. Before proceeding with this analysis we ascertained whether the flies maintained comparable velocity amplitudes across all conditions *(conditions* refers to the presence or absence of one of the four possible distractors). This was done in order to avoid any bias due to variations in velocity determined by the experimental conditions. We found no evidence for differences in velocity amplitude across conditions (Fig. 4A, B). Next, a series of LME models were fitted to the fly trajectory data (first two seconds following the presentation of the distractor) in order to obtain the best-fit model explaining the spatial orientation of flies as a function of time. The best fitting model (the one having the lowest BIC) was the following:

**Fig. 4.**
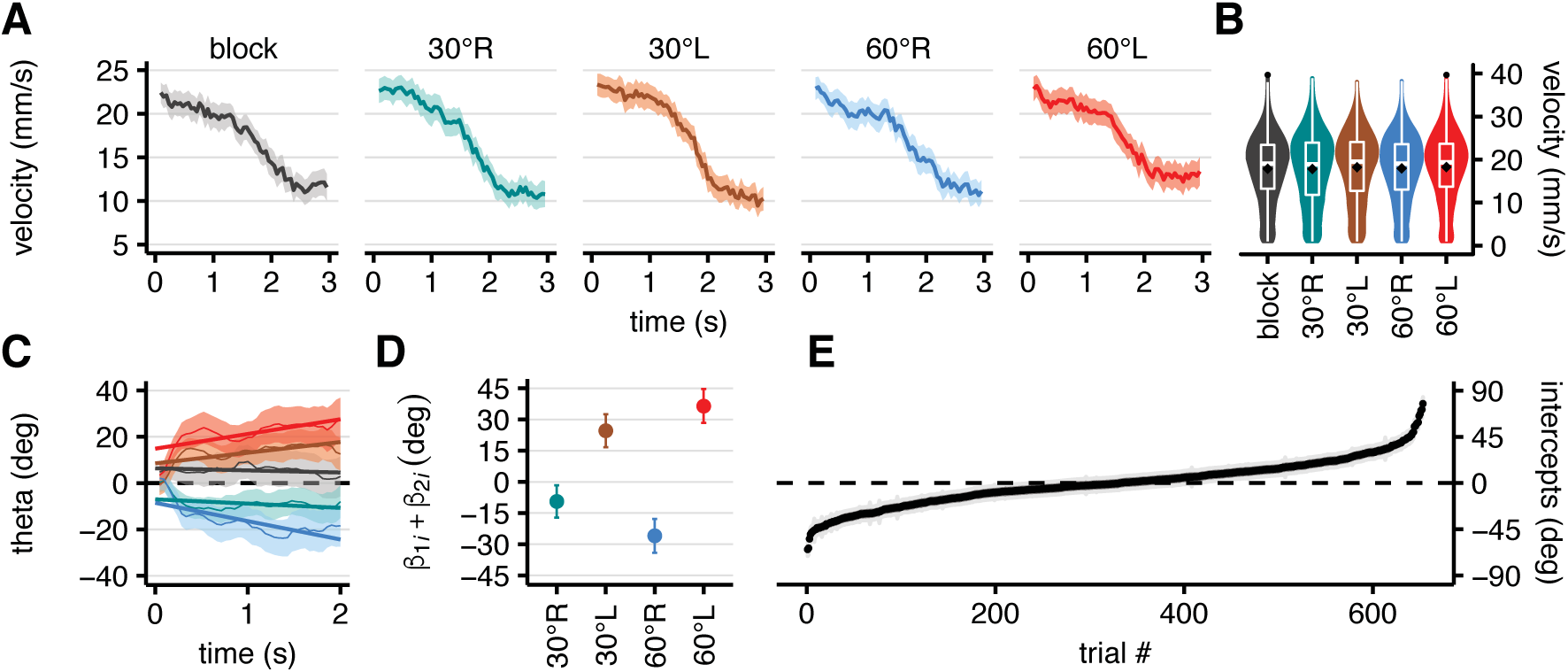
Plots of orientations. (A) Velocity profiles from t = 0 s (crossing of the virtual central window; see Fig. 2C) to t = 3 s in the five conditions during the task. Shaded regions represent the s.e.m. (B) Box-violin plot (i.e. box plot plus the data distribution) of the velocity values in the five conditions. One-way ANOVA provides no evidenced of differences between mean velocities among the five conditions (F_(4, 31)_ = .22, R^2^ = .53, p = .93). (C) Plot of the mean orientation (theta) from t = 0 to t = 2 s in the five conditions. Shaded regions represent s.e.m. Thick lines are regression lines for each condition. (D) Sum of the two coefficients *β_1_* and *β_2_* both referred to the condition effects (i.e. intercept and interaction with time), which allows to grasp the amount of change in orientation. (E) Random effect plot for each trial nested within flies (fly:trial). Dots represent the conditional means (also known as BLUPs, Best Linear Unbiased Predictions) while the shaded region (grey) corresponds to the standard deviations. In all images of the panel, the color-coding is as previously described (see Fig. 3).

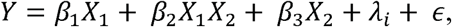

*Y* represents the predicted orientation, *β*_1_and *β*_3_ are the intercepts of regressions represented by the condition and time variables, respectively, while *β*_2_ is the slope that represents the interaction between conditions and time. Finally, *λ_i_* represents the random effect which results in variation of the regression intercepts among trials within flies, while *ε* represents the error component. At a first glance, the linear regressions relating to the fixed effects (i.e. the orientation of flies in relation to the experimental condition) show that flies tend to orient consistently towards the distractor (Fig. 4C) suggesting an influence of the distractor on the orientation of the flies. However, as the data in Fig. 4C also suggest, flies did not tend to turn fully towards the distractor. This can be more clearly appreciated by evaluating a summary-measure of the orientation predicted by the LME model, that is, the sum of the two coefficients *β*_1_ and *β*_2_ which in the LME model both refer to the experimental condition effects (i.e. condition and condition-time interaction, respectively). This provides a more direct and global representation of the change in orientation of flies following the presentation of the distractors – showing that the orientation of flies does not precisely match the expected orientation based on the position of the distractor (Fig. 4D). Rather, the model predicts that the orientation of flies, following distractor presentation, is intermediate between the orientation of the original trajectory and that of the distractor-influenced trajectory. Fig. 4E represents the distribution of the random effects. Given the significant length of each experimental session (i.e. approx. 30 min) we also evaluated the possibility that flies could show signs of fatigue across trials, which in turn might affect the re-orientation behaviour. Inspection of the average velocity profiles with respect to time for each trial does not suggest the onset of fatigue, which would presumably result in a systematic decrease in velocity as a function of time (supplementary material Fig. S1).

### Spatial trajectories

Considering the idea that distractors could act on flies through a novelty effect, as a measure of the flies’ commitment to move towards the stimuli we explored the displacement of flies along the x-axis at mid-path (i.e. after the flies had travelled 24 mm following the presentation of the distractor). We hypothesized that, given the premise, there might be a reduction in the shift of the flies’ trajectory towards the new target whenever the target presented was (randomly) preceded by one of the same kind (i.e. on the same side and at the same angle, in which case it would not be interpreted by the fly as a novel stimulus). Interestingly, a tendency consistent with this idea could in fact be observed (Fig. 5A). As a corollary, distinct left and right shifts (depending on the type of distractor presented) were evident at the end of the paths (Fig. 5B), meaning that flies not only re-oriented toward the distractor but that in so doing, they also committed to a new path (for individual tracks see supplementary material Fig. S2). In order to obtain a model of the flies’ trajectories, which would provide an objective and quantitative evaluation of the strength and the extent of the tendency of flies to shift their trajectories towards the distractors, we tested seven LME models (Table 3). To this end we considered only trajectories at least 45 mm long, (which corresponds to the radius of the surface of the arena effectively explorable by flies), were considered. The best LME (i.e. the one with the lowest BIC), LME_6, was a very parsimonious model consisting of only one *β*_1_ interaction parameter (representing the interaction between distance (d) and ‘distractor’ condition as a fixed effect, d:condition, Fig. 5C) in addition to a stochastic variation in the intercept among trials within flies (fly:trial, Fig. 5D) as a random effect:

**Table 3.**
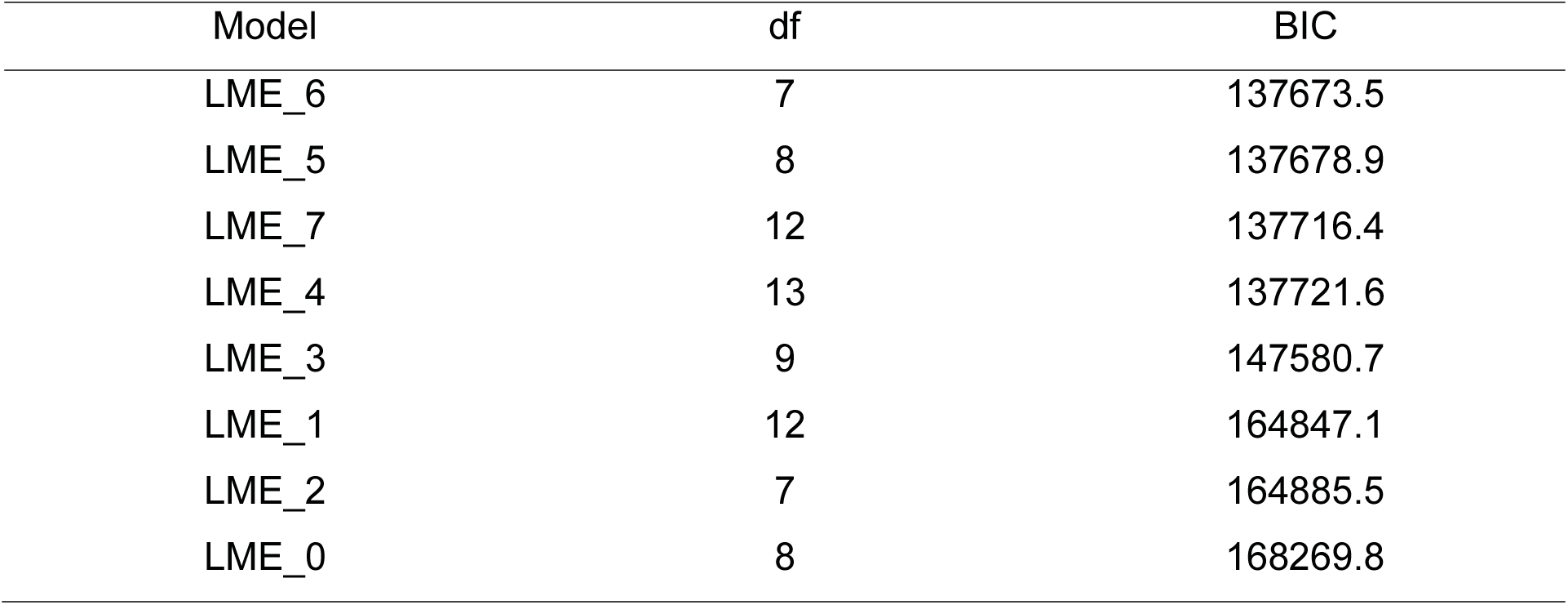
BIC of LMEs

**Fig. 5.**
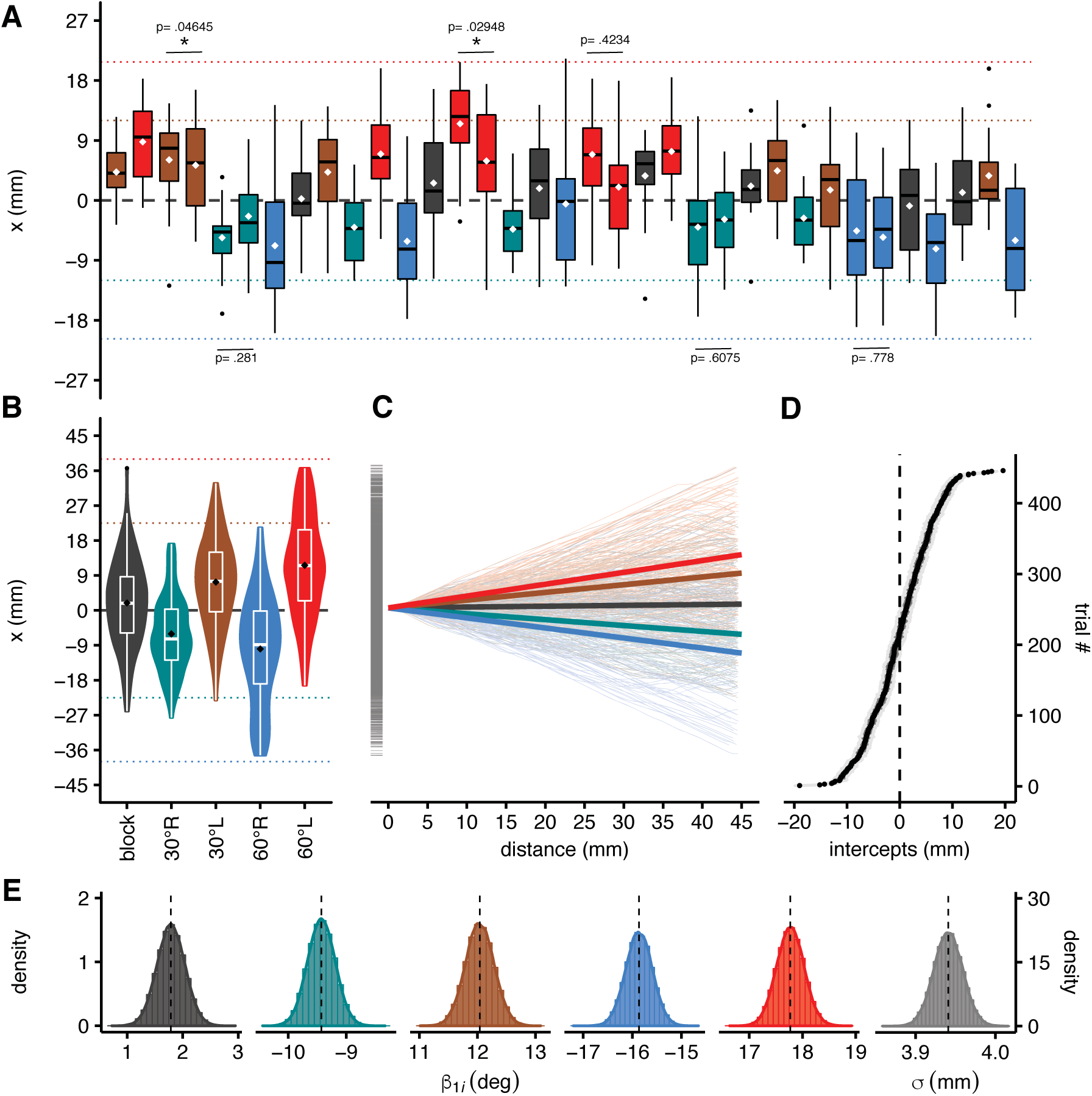
Plots of trajectories. (A) Box and whiskers plot of the displacement of flies along the x-axis at mid-path (i.e. after the flies had travelled 24 mm following the presentation of the distractor). Each trial corresponds to a specific condition and is presented across time from left to right. This allows to appreciate the horizontal shift of the trajectories, at midway, between trials. This graph shows two measures of the central tendency, the median as a black horizontal line inside the box plot and the mean represented by a white squared dot. The vertical extension of each box represents the interquartile range (IQ). The whiskers extending from each box represent the extension of the data (i.e. max. and min. of the data within 1.5 times the IQ), while isolated black dots represent outliers. It can be observed that when the same distractor is immediately re-presented (which can only occur occasionally, due to the randomness of distractor presentation), the shift along the x-axis is usually smaller than the shift observed when the distractor is presented for the first time or has not been presented recently. Only on two occasions out of the six, did the mean displacement values between two successive presentations of the same distractor differ significantly (p = .04645 and p = .02948). Statistical comparisons were done using the non-parametric Mann-Whitney-Wilcoxon test. (B) Box-violin plot (i.e. box plot plus the data distribution) of the displacement of flies along the x-axis for each condition when flies have travelled for 45 mm (i.e. along the axis connecting the two fixed stimuli) by condition. (C) Plot of the regression lines (thicker lines) for each condition with the intercept fixed at x = 0 for all trajectories (thinner lines). (D) Plot of conditional modes of the random effects of the LME_6 model. Dots represent the conditional means (also known as BLUPs, Best Linear Unbiased Predictions) while the shaded region (grey) corresponds to the standard deviations. This represents the difference between the average predicted response for a given condition and the response predicted for a particular individual. (E) Approximate density profile of the probability density function for the sampling distribution for each parameter. The six distributions show the likelihoods of the five interaction parameters (between distance and condition), with σ representing the residual standard deviation. In all images of the panel the color-coding is as previously described (see Fig. 3).

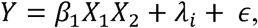

In this case, *Y* represents the displacement of flies along the x-axis. This implies that the best model represents effects as changes in the slope of the fitted line (which represents the interaction), according to the ‘distractor’ condition (Table 4). An estimate of the Confidence Intervals (CI) of the interaction parameters (Table 5) shows that none of them overlap which, in the classic frequentist perspective, implies a statistically significant difference between the effects of different conditions. The predictor (*β*_1_) can be converted into an angular measure by means of a simple trigonometric conversion:

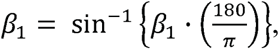

and in this way it is possible to highlight the direction of flies predicted by the model (Fig. 5E). As already seen in the case of the fly body orientations, albeit to a lesser extent, the trajectories of the flies also shifted coherently with the distractor position (i.e. the greater the angle of the distractor with respect to the original trajectory, the farther the flies’ path shifted in the direction of the distractor). None of the trajectories’ regression per condition seems to predict an angulation (with respect to the fly) superimposable to the real angle subtended for both the ‘block’ and the ‘distractor’ conditions. Flies ended between the two but closer to the original target, with a little difference between the 30 and the 60 deg conditions.

**Table 4.** Coefficients of the LME_6

| Parameter | Estimate | Std. Error | t value |
| --- | --- | --- | --- |
| d:condition block | 0.0312009 | 0.0044081 | 7.078053 |
| d:condition 30°R | -0.1638399 | 0.0040951 | -40.008641 |
| d:condition 30°L | 0.2085999 | 0.0042346 | 49.261415 |
| d:condition 60°R | -0.2733438 | 0.0045307 | -60.331208 |
| d:condition 60°L | 0.3052126 | 0.0042944 | 71.072942 |

**Table 5.** Estimated C.I. of parameters

| Parameter | 2.5% | 97.5% |
| --- | --- | --- |
| sd_(Intercept)\|fly:trial ($\sigma_1$) | 5.7658491 | 6.5809188 |
| $\sigma$ | 3.9055042 | 3.9762879 |
| d:condition block | 0.0225617 | 0.0398401 |
| d:condition 30°R | -0.1718659 | -0.1558140 |
| d:condition 30°L | 0.2003008 | 0.2168990 |
| d:condition 60°R | -0.2822242 | -0.2644633 |
| d:condition 60°L | 0.2967963 | 0.3136289 |

### Bayesian trajectories model

The BF analysis highlighted a less parsimonious model with respect to the one which was selected using the frequentist approach:

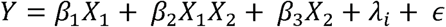

This model, in addition to a *β*_2_ interaction term (d:condition), also presented the *β*_1_and *β*_3_ parameters, which individually represent the effects of experimental condition and distance, respectively. In the case of this model, the distribution of parameters and the goodness of fit were evaluated (i.e. the standard error of residuals and the R-squared were estimated), in order to assess the goodness of the model (Table 6). In this case, a “confidence interval” was computed, based on the Highest Posterior Interval (HPI), using the R package *TeachingDemos* (Snow, 2016) (Table 7). In practice, all points in an HPI region have a higher posterior density than points outside the region. For this reason HPI is also called Highest Density Interval (HDI). Notwithstanding the slightly more complex model produced by the BF analysis, this model provided essentially the same general explanation for the experimental data as the LME model. Also in this case, none of the parameters bound to the ‘experimental condition’ variable showed any overlap in the predicted values in terms of HPI (Fig. 6A), suggesting that the distractors produced significant deviations of the flies’ trajectories both in terms of shift and slope. The *β*_3_ parameter (i.e. distance) showed a non-significant shift in the intercept of the regressions (Fig. 6B), while the *λ*_*i*_ random effect due to the variation between flies appeared minimal (Fig. 6C). This analysis confirmed that flies respond to distractors by shifting their locomotor trajectories essentially in accordance with the location of the distractor, albeit not proportionately. In fact, with distractors presented at 30 deg flies tended to adopt a heading of 10 deg, while with distractors presented at 60 deg flies adopted a heading of 16 deg.

**Table 6.** Model goodness of fit

| Residual-se | R-squared |
| --- | --- |
| 7.199331 | 0.4120791 |

**Table 7.** HPI of parameters

| Parameter | 2.5% | 97.5% |
| --- | --- | --- |
| distance | 0.0073964 | 0.0209496 |
| condition block | -0.0433319 | 0.3353007 |
| condition 30°R | -4.2541136 | -3.8968358 |
| condition 30°L | 3.8712083 | 4.2320447 |
| condition 60°R | -6.9712379 | -6.5951677 |
| condition 60°L | 6.4814631 | 6.8356365 |
| d:condition block | -0.0303948 | -0.0031744 |
| d:condition 30°R | -0.1868404 | -0.1609655 |
| d:condition 30°L | 0.1671820 | 0.1942019 |
| d:condition 60°R | -0.2949580 | -0.2669707 |
| d:condition 60°L | 0.2775440 | 0.3045376 |
| $\sigma^2$ | 50.9114193 | 52.7319502 |

**Fig. 6.**
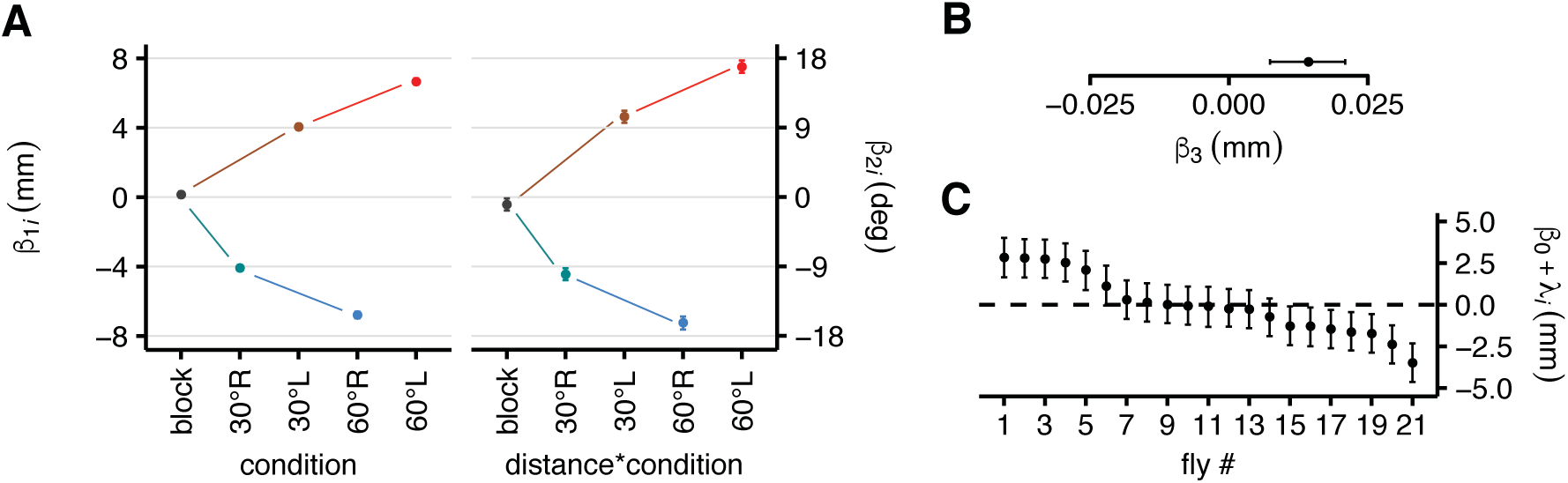
HPI plot of parameters. (A) Fixed effects of condition parameter (on the left) and interaction parameter (on the right) between distance and condition with their 97.5% Highest Posterior Intervals (HPI). (B) Fixed effects of distance parameters with their 97.5% HPI. (C) Random effects plot of the model represented for each fly. Colours encode conditions as previously described (see Fig. 3).

### Kinematics indices

The ‘partial attraction’ effect determined by the appearance of distractors led us to hypothesise that perhaps a high number of the trajectories used in the model construction and analysis were trajectories of flies which remained on the original straight path (i.e. which essentially did not respond to the distractor), impacting on the leverage of the model. Indeed, the raw distribution of the position of flies within the arena shows peaks which are consistent with the position of the original target (supplementary material Fig. S3). In order to clarify this issue we considered separately two situations: (i) the trials in which the distractor determined only a slight shift in the fly’s trajectory in that direction, with the fly essentially maintaining the direction towards the original target (type 1); (ii) the trials in which the presence of the distractor determined a dramatic change in trajectory, with the fly abandoning the original direction in favour of the one dictated by the distractor (type 2). Type 2 trajectories were selected by considering a shift of at least 9 mm from an ideal straight line – orthogonal to the original target – at the time the individual crossed the middle of the path. This arbitrary procedure did not affect the balancing of the trials per condition in favour of one of the two types, maintaining a similar numerosity in the ‘block’ condition (Fig. 7A). Following this, a new parameter (i.e. shift) was introduced in the LME model as a third component of the interaction between distance and condition, thus increasing the values of the predictors (Fig. 7B). This kind of manipulation allowed us to investigate possible changes in kinematics following the appearance of the distractor. During the first 21 frames (i.e. 1 s), the flies executed a fast turn in response to the distractor (Fig. 7C). In particular, around 250 ms the type 1 flies began to perform a body saccade in the contrary direction, while type 2 flies continued to maintain an orientation which was coherent with the distractor position (Fig. 7D). These fast turns did not affect the final trajectories of the flies (Fig. 7E).

**Fig. 7.**
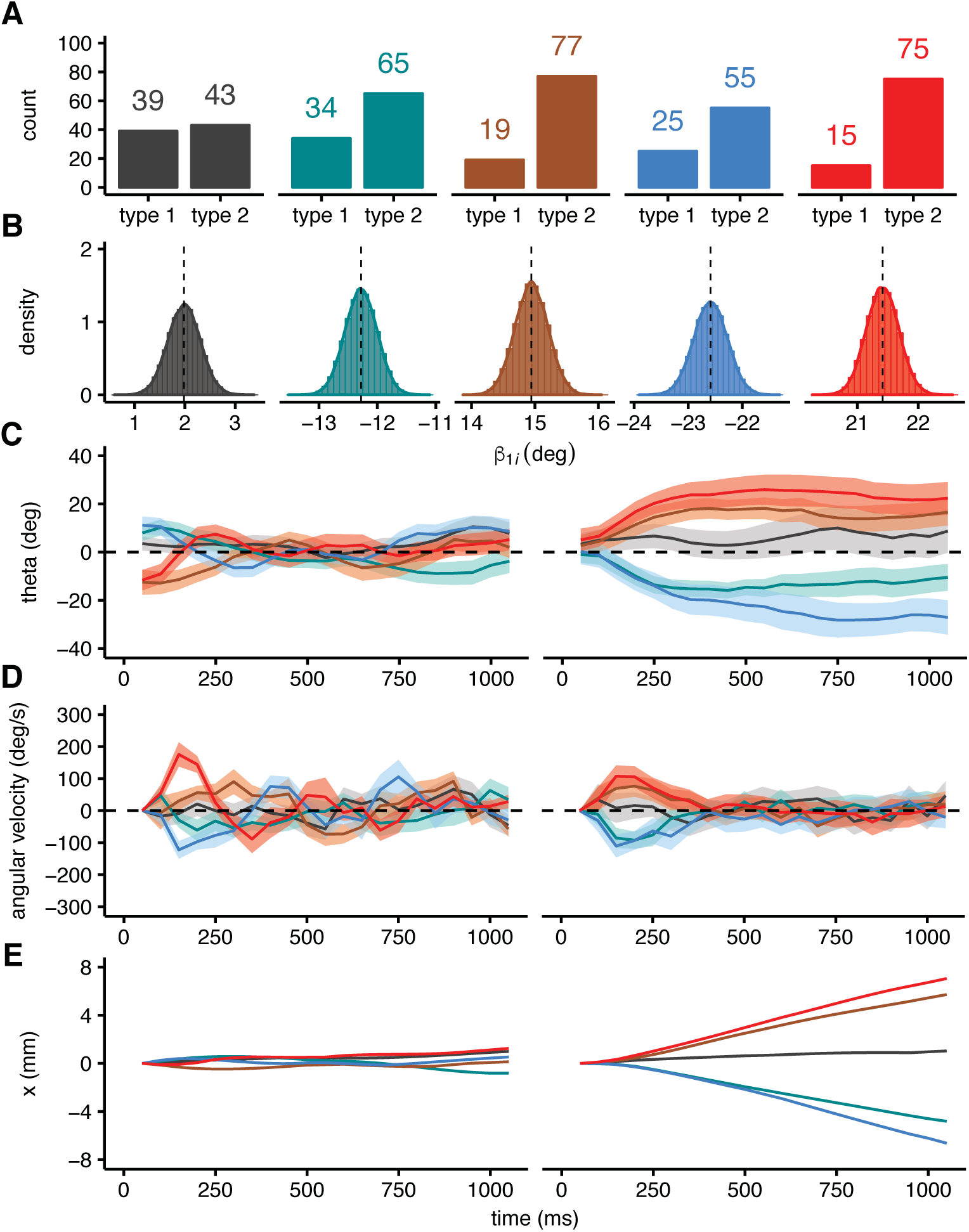
Trajectories split. (A) Count of the trials falling into the two types of trajectories by condition. Type 1 represents the trajectories in which the shift of the flies was at least 9 mm from an ideal straight line uniting the two Buridan stimuli when flies reached the middle of the path, while type 2 represents the trajectories for which the shift was less than 9 mm. (B) Approximate density profile of the probability density function for the sampling distribution for the five conditions. Distributions show the likelihoods of the interaction parameters (between distance, condition and type). (C) Mean orientation of the flies during the first second by condition in the two types of trajectories. On the left are shown type 1 trajectories while on the right type 2 trajectories. The shaded region represents the s.e.m. (D) Mean of the angular velocity of the flies during the first 1000 milliseconds by condition in the two types of trajectories. Type 1 on the left, type 2 on the right. The shaded region represents the s.e.m. (E) Regression lines of the trajectories with the LOWESS (LOcally WEighted Scatterplot Smoothing) method during the first 1000 milliseconds by condition for the two types of trajectories. Type 1 on the left, type 2 on the right. In all images of the panel the color-coding is as previously described (see Fig. 3).

## DISCUSSION

The primary aim of this research was to evaluate in what way the abrupt presentation of different distracting visual stimuli to fruit flies which are already engaged in locomotion (walking) towards a pre-existing visual target, would influence the original locomotion action. Our results indicate that, following the presentation of a distractor, flies oriented their bodies according to a vector positioned midway between the original target and the distractor. Following the initial body orientation, flies then engaged in locomotion by committing to a new trajectory, essentially in one of two ways: (i) the presence of the distractor produced a slight perturbation in the original trajectory, but the ensuing movement then tended to proceed in the direction of the original target; (ii) the presence of the distractor determined the insurgence of an alternative motor program, which had the power to override the original one, leading to a dramatic change in the direction of the flies’ motion.

### Buridan with light stripes

First and foremost some considerations concerning the use of the type of stimuli used here are in order. By using a tethered flight simulator, it has been demonstrated that flies are usually attracted towards long vertical bright or dark stripes, as an ethological reflex which guides flies towards elements resembling vegetative perches (Maimon et al., 2008). Here we describe for the first time the free walking behaviour of flies consisting of recurrent orientation inversions (i.e. alternation between fixation and anti-fixation) between two diametrically opposed vertical bright stripes on a dark background. Pioneer studies had shown that recurrent inversion is maximized with vertical black stripes on a bright background (Bülthoff et al., 1982) and had considered the opposite contrast as a repellent configuration for flies (Heisenberg and Wolf, 1979). Notwithstanding these earlier observations, we observed a strong fixation response toward bright stripes in freely walking flies consistent with more recent studies using tethered flying flies (Reiser and Dickinson, 2008; Maimon et al., 2008). We are tempted to exclude that the recurrent direction inversions shown by the flies in our case was due to anti-fixation, because when presented with the distractor stripes flies were attracted to and maintained the distractor in front of them (suggesting fixation). Although the functional distinction between flying and walking paradigms, as well as differences in the experimental protocols, such as wing clipping (McEwen, 1918; Gorostiza et al., 2016), might be at the basis of these contrasting findings, it is difficult to draw a coherent explanatory picture, and the exact reason for the discrepancies remains as yet unknown. Rather, it is possible that the intensity of the light used may have played a role in determining the discrepancies concerned with anti-fixation behaviour of the flies, since in the case of LED displays (such as those used in the present study) the maximum luminance reachable is 72 cd m^−2^ (cd m^−2^ = lux) (Reiser and Dickinson, 2008), while in the setups used in previous studies the luminance ranged between 300 and 1910 cd m^−2^ (Götz and Wenking, 1973; Bülthoff et al., 1982; Virsik and Reichardt, 1976), which is at least 4 times higher. This suggests that long vertical bars with high light intensities lead to avoidance, while long vertical bars of lower brightness (i.e. in the region of 72 lx) would represent an attracting stimulus, possibly because under these conditions the bar appears similar to the reflectance of natural vegetation posts. This hypothesis seems to be corroborated by a report of Heisenberg and Wolf (1984), in which a grey background makes bright stripes as attractive as black stripes on a white background, while bright stripes on a black background produce anti-fixation behaviour (Heisenberg and Wolf, 1984).

### Novelty effect

One aspect of the present results suggests that in our experimental paradigm the appearance of the visual distractor stimuli produced a novelty effect in the flies. In particular this was manifested by the re-orienting behaviour of the flies immediately following the appearance of the distractors. A similar effect has been reported for tethered flying flies which showed a preference for a previously uncued side of the arena when faced with bilateral stimuli (Shiozaki and Kazama, 2017). In neural terms, it has been suggested that the *Drosophila* EB ring neurons (R4), are involved in determining the slow turning tendency (i.e. body re-orientation) associated with this kind of visual experience. Silencing of those neurons abolishes the innate behaviour for preferential orientation toward novel stimuli (i.e. previously uncued sides) (Shiozaki and Kazama, 2017).

In another study using calcium imaging, the authors found that visual responses in ring neurons are suppressed when competing stimuli are present in the contralateral visual field (Sun et al., 2017). In this respect, contralateral suppression is hypothesized to act as a mechanism for location-based stimulus selection by reducing the responses of ipsilateral stimuli in the presence of a second stimulus. Furthermore, this suppressive effect appears to dependent upon short-term stimulus history, specifically, ring neurons baseline activity showed a rebound after contralateral suppression, a phenomenon which could be involved in modulating the flies’ subsequent visual responses to both ipsilateral and contralateral stimuli (Sun et al., 2017). Such evidence could partially explain our results, at least in terms of the novelty represented by the distractor.

The EB ring neurons – which innervate four concentric rings within the EB – appear to be retinotopically modulated by visual patterns but not by locomotor states (Seelig and Jayaraman, 2013). These neurons are possibly upstream from the EB wedge neurons, and convey visual information to the integrator layer. In fact, some of these neurons (R4d and R3) have been implicated in visual working memory (Neuser et al., 2008) and others (R4 and R1) in space-learning linked to visual patterns (Ofstad et al., 2011) without affecting locomotor activity. Our findings add to this literature by showing that flies are attracted by a novel visual stimulus and that the attraction is manifested not only through a re-orienting of the body, but also by the ensuing commitment of the individual to a new locomotor path.

### Reactive turning tendency

Our data are consistent with the ‘reactive turning tendency’ described by Horn and Wehner (1975), who noted that flies preferred to orient toward a position midway between two vertical stripes placed at an angular distance less than 60 deg (Horn and Wehner, 1975). In our paradigm, the sudden appearance of the distractor added a ‘turning tendency’ of the body to the one already engaged by shifting the internal compass needle toward the distractor. Differently from what reported by Horn and Wehner (1975), we observed that the trajectories did not lie midway between the original stimulus and the distractor, but instead remained closer to the former, meaning that the original stimulus had acquired the status of a stronger landmark. In our opinion, this behaviour might be the indication of a well-established motor program which is relatively ‘impermeable’ to the possible perturbation determined by the appearance of the distractor. This is in line with the observation that the E-PG neurons show a persistent activity maintaining the compass needle information even when the animal is in total darkness (Seelig and Jayaraman, 2015). The activity of such neurons remains linked to the position of a single vertical stripe even in the presence of a second identical stripe. Furthermore, the activity of such neurons does not always shift instantaneously following the abrupt displacement of a single visual target (Seelig and Jayaraman, 2015). Therefore, it would seem that the accomplishment of a coherent motor program requires locking on to a target.

### Selection for action via inhibition

We were interested in understanding how flies detected and reacted to an abrupt distraction during the execution of a motor program. According to our original hypothesis we expected the distractor stimuli to determine an inhibitory or attracting behaviour acting upon already programmed trajectories, similarly to the interference effect observed in human and non-human primates under analogous circumstances (Tipper et al., 1998; Sartori et al., 2014; Bulgheroni et al., 2017). In these studies, participants were instructed to initiate a reaching movement after two stimuli (a target and a distractor) were presented. When the investigators compared a condition in which the target was presented alone with that in which there was a distractor acting as an alternative potential target, they found that the reaching path was affected in the latter case with the arm trajectory deviating away from or nearer to the distractor. This was observed even with regard to distractor objects that were unlikely obstacles to the reaching action. As those objects are also included in the initial processing of the whole context in which the action will be carried out, the motor program appropriate to reaching them is also produced in parallel, thus producing trajectory changes (Tipper et al., 1992; Tipper et al., 1997; Bulgheroni et al., 2017). This effect has been explained in terms of selective attention mechanisms mediating the selection of objects for action, with a specific mechanism acting to inhibit competing internal representations of distractor objects (Tipper, 1985; Tipper et al., 1992; Meegan and Tipper, 1998). Put simply, the effects caused by the presence of nearby objects seem to reflect inhibitory mechanisms. When the target is identified, the reaching movement towards the non-target is inhibited. But because there is an overlap between the target and the non-target(s), the act of reaching towards the target is affected by this non-target inhibition. Another crucial aspect of this model is that the amount of inhibition might be determined by the levels of activation of perceptual inputs. That is, inhibition is reactive such that its level is determined by the relative salience of the distractor. Thus distractors causing greater levels of neural excitation receive greater levels of inhibitory feedback. In the present circumstances our flies exhibited two kinds of behaviour in response to the distractor. The majority of flies fully espoused the new path dictated by the distractor. The remaining flies, maintained the original path with only a slight deviation toward the distractor. In both cases the flies acknowledged the presence of the distractor by making a fast saccade movement toward it within the first 250 ms from the onset of locomotion. This early fast saccade response could rely on the optomotor system, via the horizontal system neurons (HS; Bahl et al., 2013; Kim et al., 2015; Fujiwara et al., 2016). Nonetheless, to explain the present results our preferred idea is that inhibitory processes in *Drosophila melanogaster* occur at the level of the neuroanatomical structures involved in heading behaviour (Seelig and Jayaraman, 2015). This implies the involvement of the CC and in particular of the EB, with specific reference to the role played by dopamine in releasing and inhibiting motor programs. Similarly to what occurs in the mammalian brain (Grillner and Robertson, 2016), the signal involved in starting and halting an action sequence could be based on phasic dopamine release onto the EB in a manner similar to what is observed in the case of the nigrostriatal circuit of mice (Jin and Costa, 2010). The quantitative modulation of dopamine, via different receptors and/or perhaps through different types of neurons (Green et al., 2017), could engage and disengage the action programs, by respectively strengthening or weakening the inhibitory process. A high level of phasic release might enhance the specificity of action selection processes and movement initiation, while tonic release might inhibit the modules for action. This double mechanism would facilitate the emergence of motor responses from a repertoire of possible actions in order to readily cope with the sensory inputs determined by environmental variations. Fiore and collaborators (2015) suggest that a phasic dopamine release would allow the system to change the strength of the connections between sensory inputs and the EB, thus affecting the probability that the related motor action would be selected again. Conversely, a tonic release would not alter the connections’ strength but would make the global system more stable (i.e. maintenance of selection) or unstable (i.e. sensitive to changes) depending on the receptor type involved (Fiore et al., 2015). However, it remains unclear how the system would differently weigh opposing pathways in order to regulate action selection. In this respect, our paradigm might provide a novel theoretical and methodological territory within which to classify and distinguish different mechanisms concerned with action selection in flies. Further research, considering the manipulation of the neuroanatomical circuit discussed above, is needed in order to dissect the neural mechanism underlying the action selection.

## ACKNOWLEDGMENTS

We thank Fabian Feiguin (Neurobiology Lab, International Centre for Genetic Engineering and Biotechnology) for the Oregon-R fly stock, Paola Cisotto and Fortunato Piron for technical assistance.

